# Medioapical contractile pulses coordinated between cells regulate *Drosophila* eye morphogenesis

**DOI:** 10.1101/2023.03.17.529936

**Authors:** Christian Rosa Birriel, Jacob Malin, Victor Hatini

## Abstract

Lattice cells (LCs) in the developing *Drosophila* retina constantly move and change shape before attaining final forms. Previously we showed that repeated contraction and expansion of apical cell contacts affect these dynamics. Here we describe a second contributing factor, the assembly of a medioapical actomyosin ring composed of nodes linked by filaments that attract each other, fuse, and contract the LCs’ apical area. This medioapical actomyosin network is dependent on Rho1 and its known effectors. Apical cell area contraction alternates with relaxation, generating pulsatile changes in apical cell area. Strikingly, cycles of contraction and relaxation of cell area are reciprocally synchronized between adjacent LCs. Further, in a genetic screen, we identified RhoGEF2 as an activator of these Rho1 functions and RhoGAP71E/C-GAP as an inhibitor. Thus, Rho1 signaling regulates pulsatile medioapical actomyosin contraction exerting force on neighboring cells, coordinating cell behavior across the epithelium. This ultimately serves to control cell shape and maintain tissue integrity during epithelial morphogenesis of the retina.

**Short summary:** Rosa et. al. describe a Rho1-dependent pulsatile medioapical actomyosin network that couples neighboring retina lattice cells biomechanically, creating an adaptive supracellular actomyosin network that coordinates the mechanical behavior of neighboring cells and regulates cell shape and tissue integrity.

## INTRODUCTION

Morphogenesis of the *Drosophila* retina involves changes in cell shape, size, and junction length that seem largely transient. Through this process, which unfolds over tens of hours, epithelial cells achieve their precise number, shape and arrangement. The result is a tessellated structure composed of approximately 800 nearly identical ommatidia (Cagan and Ready, 1989). This process is a model for understanding morphogenesis of differentiated cells forming a mature organ. Each ommatidium consists of four central cone cells, surrounded by two large semi-circular primary (1°) cells. These are surrounded by lattice cells (LCs) and mechanosensory bristles (Fig. 1A). Of these, the LCs exhibit a particularly intricate developmental sequence in which they intercalate to arrange in a single file between ommatidia, extra LCs are pruned, and remaining cells undergo dramatic shape changes according to their location (Cagan, 2009; Carthew, 2007; Johnson, 2021).

**Figure 1:**
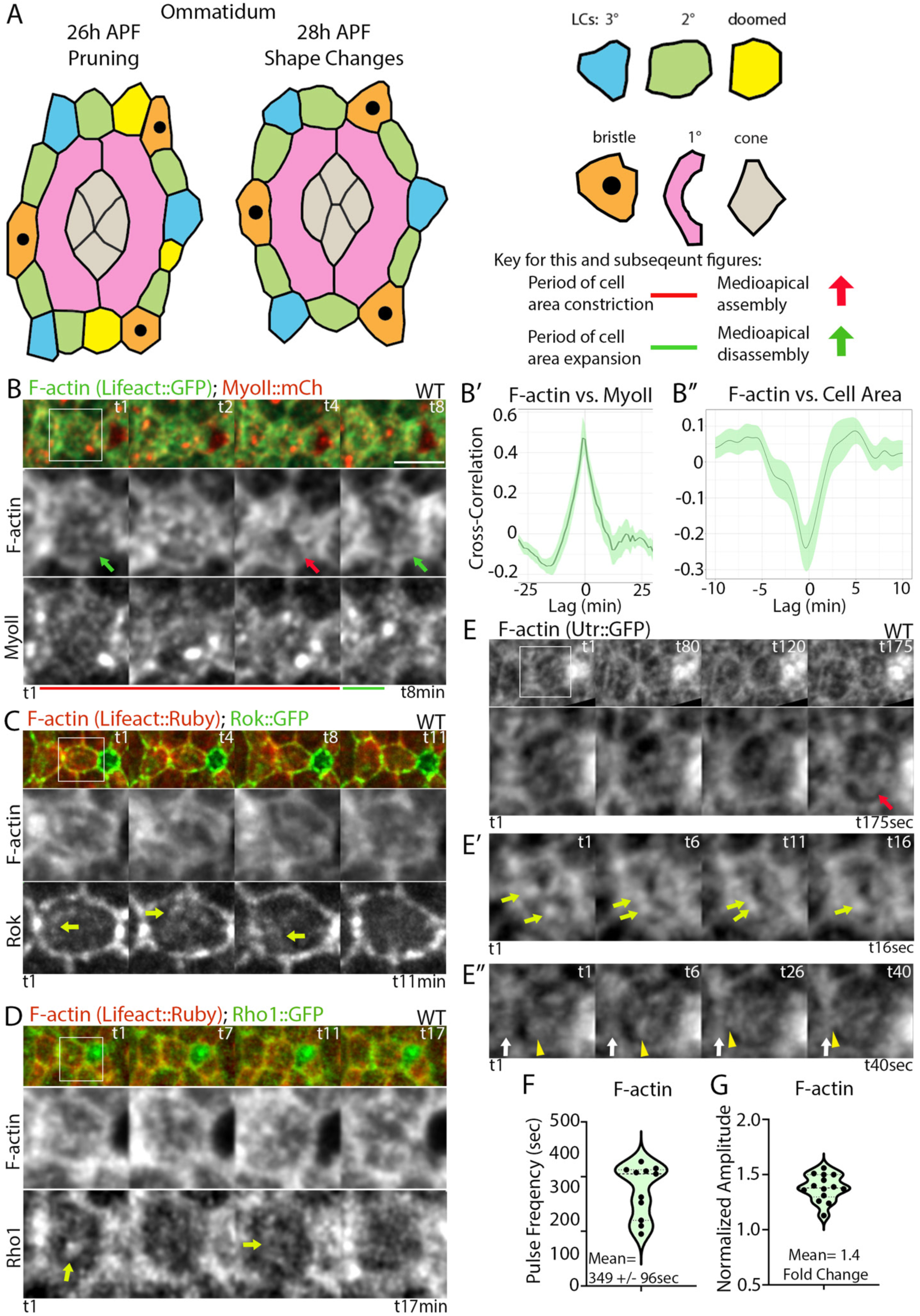
A dynamic medioapical actomyosin network correlates with cell area fluctuations during epithelial remodeling. (A) Left: Cartoon of *Drosophila* ommatidium during pruning and shape changes. Right: Key for cell types and mechanical and cytoskeletal dynamics in this and subsequent images. (B) Medioapical actomyosin dynamics in LCs during cell shape changes. Top: Snapshots of a lattice edge highlight the assembly (red arrow) and disassembly (green arrow) of a medioapical actomyosin ring. Bottom: Zoomed-in images of F-actin and MyoII dynamics. Red bar: Assembly of a medioapical ring; Green bar: Ring disassembly. (B’-B’’) Time-shifted Pearson’s correlation charts. Solid line, mean R-value at each time shift; Green band, standard error of the mean, here and in subsequent figures. (B’) Medioapical F-actin and MyoII accumulation positively correlate. (B’’) F-actin and cell area negatively correlate. (C) F-actin and Rok dynamics. Top: Snapshots of a lattice edge. Bottom: Zoomed-in views of F-actin and Rok dynamics in a LC. Yellow arrows: Medioapical foci of Rok accumulation. (D) Rho1. Top: Snapshots of a lattice edge. Bottom: Zoomed-in views of F-actin and Rho1 in a LC. Yellow arrows: foci of Rho1 accumulation. (E-E’’) Kymographs show three behaviors of the medioapical actomyosin network in LCs: (E) Assembly of a medioapical ring (red arrow) during LC contraction. (E’) Movement and fusion of F-actin nodes (arrows) during ring formation. (E’’) Movement of nodes (arrowheads) and fusion with the junctional actomyosin network (arrow) during ring disassembly. (F) Medioapical F-actin pulse frequency in WT 2° LCs is approximately 6 minutes (349±96 sec, N=11). (G) On average, medioapical F-actin levels are 1.4-fold higher at their peak than at their trough levels. Scale bar in this and subsequent figures is 5 µm.

The initial arrangement of cells in ommatidia is controlled by differential adhesion (Bao and Cagan, 2005; Bao et al., 2010; Hayashi and Carthew, 2004). After the intercalation of LCs, certain cytoskeletal networks are preferentially associated with a subset of LCs’ contacts. Specifically, we showed that contractile junctional actomyosin networks assemble at and contract the LC-LC contacts. These contractile networks are counterbalanced by protrusive branched actin networks that assemble along the same contacts with reciprocal dynamics (Del Signore et al., 2018; Malin et al., 2022). This interplay of contractile and protrusive forces along cell contacts guides, at least in part, their movements, and prevents errors in cell shaping and arrangement.

In addition to the junctional cytoskeletal networks, a second medioapical actomyosin network is present in the developing fly retina. In 1° cells, this network assembles in MyoII foci that correlate with cell area contraction. Laser ablation of the network in 1° cells expanded their apical cell perimeter, indicating that the network applies tension on the cell surface (Blackie et al., 2020). However, the role of this network in lattice remodeling, its dynamic properties, and regulation by upstream signals remain unknown. Medioapical actomyosin networks have been observed during epithelial remodeling in several processes in *Drosophila*, including gastrulation, germband extension and dorsal closure (Azevedo et al., 2011; Blanchard et al., 2010; Fernandez-Gonzalez and Zallen, 2011; Martin et al., 2009; Rauzi et al., 2010). In those processes, medioapical actomyosin networks pulse – that is, they repeatedly assemble and disassemble. Disrupting pulsing affects cell behavior and epithelial organization. However, these early developmental processes fundamentally differ from the remodeling of the retina. They entail massive reorganization of cells that takes place over the time course of tens of minutes, compared to the precise refinement of the retina, which takes tens of hours. Furthermore, the pulsation typically displays a ratcheting characteristic, in which relaxation after constriction is only partial and the cells never revert to their original shape. In contrast, in the retina, there is no obvious ratcheting characteristic to cell-shape changes.

Cell movement and shape changes, however, are clearly critical for generating the very precise organization of the retina, because errors occur when they are perturbed (Blackie et al., 2021; Blackie et al., 2020; Del Signore et al., 2018; Johnson et al., 2008; Larson et al., 2008; Letizia et al., 2019; Malin et al., 2022; Seppa et al., 2008). Thus, a major challenge is identifying mechanisms that regulate cell dynamics and understanding their impact on force generation, force transmission and the eventual attainment of specific cell shapes.

Here we used *Drosophila* LCs to investigate the dynamic properties of the medioapical actomyosin network in apical epithelial remodeling using high-resolution live imaging and laser ablation combined with genetic tests of gene function. We asked if medioapical actomyosin networks pulse and studied their contribution to cell shape changes. We found a pulsatile medioapical actomyosin network, whose assembly promoted the transient, anisotropic contraction of LCs. Our experiments revealed the interplay between the assembly and disassembly of medioapical actomyosin networks and mechanical interactions between neighboring cells. In addition, we identified upstream regulators of Rho1, which control the assembly of the medioapical actomyosin network. While some features were similar to other developmental processes involving medioapical actomyosin networks, many aspects are unique to this late-stage developmental process. Overall, our data provide evidence that biochemical and mechanical signals that control Rho1 activation are required to rebalance forces in the epithelium, control cell shape and rearrangements and maintain tissue integrity.

## RESULTS

### A dynamic medioapical actomyosin network assembles during LC remodeling

The developing fly retina is a model for understanding how force-generating cytoskeletal networks control cell shape, cell-to-cell interactions, and tissue patterns. During lattice remodeling, the 2° LCs narrow and elongate to form the edges of the lattice while the 3° LCs compact to form alternating corners (Fig. 1A). To examine the behaviors of the medioapical actomyosin network during lattice remodeling, we live-imaged F-actin using the actin-binding domain of Utrophin tagged with GFP (Utr::GFP), together with Spaghetti Squash (Sqh), the regulatory light chain of non-muscle Myosin II (MyoII), tagged with mCherry (MyoII::mCh). In time-lapse movies, we found that F-actin and MyoII accumulated in particles that flowed from the cell surface toward the cell’s medioapical region, forming a dynamic ring-like structure (Fig. 1B, red arrow), which then disassembled (Fig. 1B, green arrows). Time-shifted Pearson’s correlation analysis confirmed that medioapical F-actin and MyoII accumulated with the same dynamics, suggesting they co-assemble to form a contractile actomyosin network (Figs. 1B’, S1A-B’, R= 0.4400). We also observed fluctuations in the LCs’ apical area (Fig. 1B) and asked if they correlated with fluctuations in actomyosin levels. We found a significant negative correlation between medioapical F-actin and LCs’ apical area (Figs. 1B’’, S1B’’, R= −0.2706). The results suggest that an increase in medioapical actomyosin levels exerts tension on the apical cell perimeter leading to a reduction in the apical cell area. This observation is consistent with a previous report showing that MyoII accumulation in medioapical foci in the 1° cells correlated with a decrease in the cells’ apical area (Blackie et al., 2020).

The Rho1 Rho GTPase activates the formin Diaphanous (Dia) to assemble linear actin filament and Rok to phosphorylate the MyoII regulatory light chain, controlling contractile actomyosin network assembly and contraction (Homem and Peifer, 2008; Magie et al., 1999; Mizuno et al., 1999; Mulinari et al., 2008; Warner and Longmore, 2009a; Warner and Longmore, 2009b). To determine if Rho1 functions medioapically, we examined the accumulation of GFP-tagged Rho1 and Rok reporters relative to F-actin marked with the F-actin-binding peptide Lifeact tagged with Ruby fluorescent protein (Lifeact::Ruby). We found that both Rho1 and Rok accumulated medioapically in dynamic particles (Fig. 1C and D, yellow arrows), suggesting that the two proteins control medioapical actomyosin dynamics.

To better understand how the medioapical actomyosin ring-like structure forms, we imaged the network at a higher temporal resolution, obtaining an image stack every 5 seconds. We found that some F-actin particles moved toward one another during ring formation and merged to form larger particles (Fig. 1E-E’; Movie 1). As the ring disassembled, some particles disappeared while others flowed toward cell edges and merged with the junctional network (Fig. 1E’’, yellow arrowheads; Movie 1). The medioapical ring assembled approximately every 6 min (Fig. 1F; 349±96 sec), with a 1.4-fold increase in F-actin levels from trough to peak in each cycle (Fig. 1G). This network behavior implies that medioapical actomyosin forms a contractile network composed of interconnected nodes that pull on one another and the cell surface in a process controlled by Rho1 and Rok.

### Medioapical actomyosin exerts tension on the apical cell surface and is mechanically adaptive

Using high-resolution Airyscan confocal imaging, we examined the structure of the medioapical actomyosin network at higher spatial resolution in time-lapse movies. We identified nodes that form the medioapical ring-like structure and filamentous structures between nodes and between nodes and the cell surface (Fig. 2A, left panel, yellow arrowheads). Some F-actin filaments connected to nodes were also linked to LC-LC contacts, which are sites that dynamically accumulate F-actin (Fig. 2A, right panel, yellow arrowheads). This architecture suggested that the network exerts tension on the cell surface and that LCs are mechanically linked through a dynamic supracellular medioapical actomyosin network anchored, in part, at LC-LC contacts.

**Figure 2:**
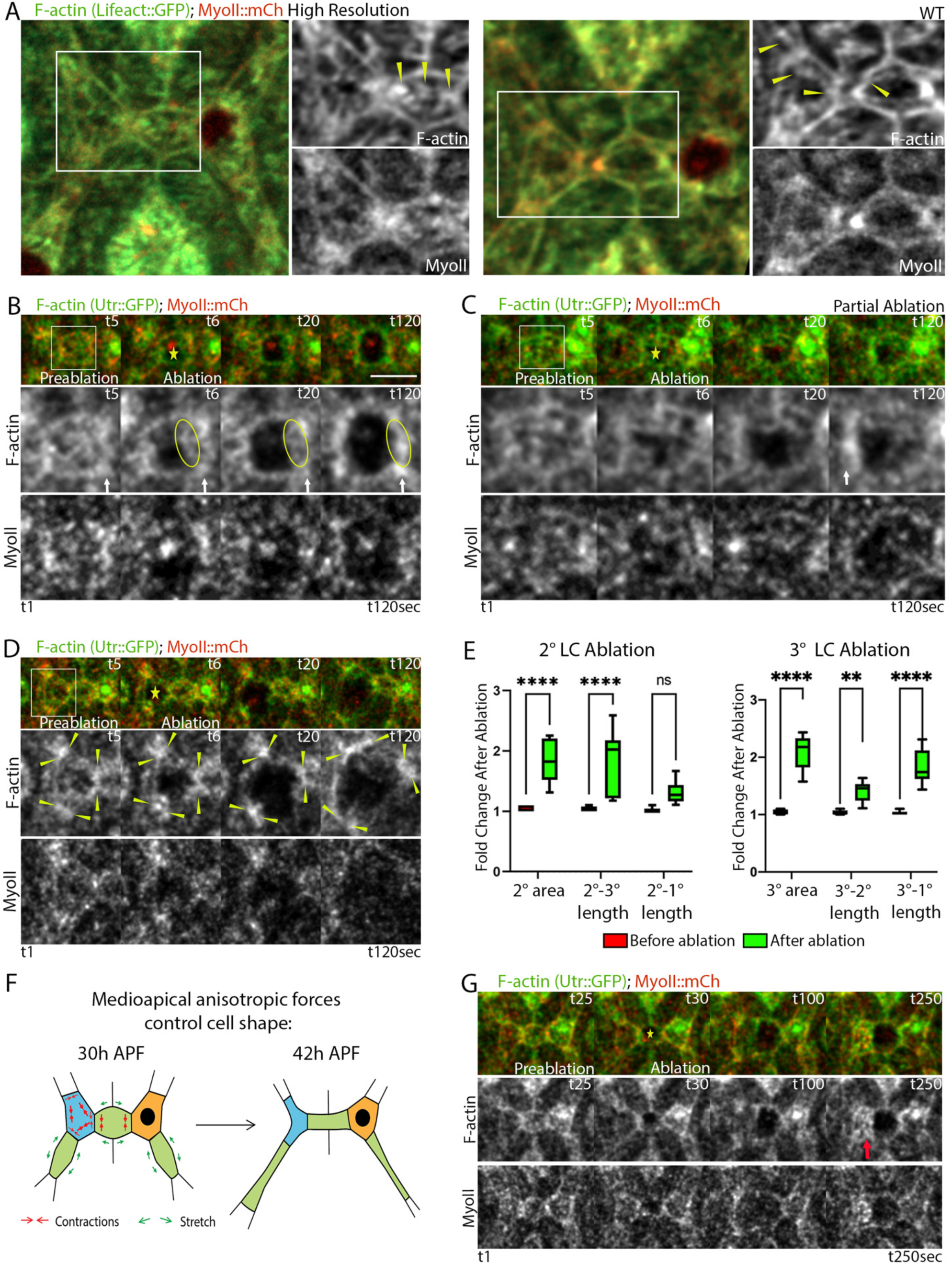
The medioapical actomyosin network holds tension and mechanically links neighboring LCs. (A) High-resolution Airyscan images of F-actin and MyoII from a time-lapse movie. Note medioapical actomyosin ring in 2° and 3° LCs. Medioapical actomyosin nodes connect to cell contacts through filamentous structures (yellow arrowheads). (B-E) Targeted ablations (marked with a yellow star) of medioapical actomyosin in LCs induced a rapid recoil of medioapical actomyosin and apical cell area relaxation. (B) Medioapical actomyosin ablation in 2° LCs induced preferential expansion of the 2°-3° contact and actomyosin flow (marked by the ellipse) toward LC-LC contacts (marked by the white arrow), concomitant with apical area relaxation. Top: Snapshots of a lattice edge from a time-lapse movie; Bottom: Zoomed-in views of F-actin and MyoII. (C) Partial medioapical actomyosin recoil to LC-LC contacts (white arrow) after ablation resulted in a weaker apical area relaxation. (D) Ablation of the medioapical actomyosin network in 3° LCs induced a strong relaxation of the apical cell area and a preferential expansion of 3°-1° contacts compared to LC-LC (3°-2°) contacts. Vertices of the ablated 3° LC are marked with yellow arrowheads. (E) Left panel: Fold change of the 2° LC area and cell-cell contacts after ablation. Two-way ANOVA was performed with Sídák’s multi-comparison to compare each data set before to after ablation. 2° LCs’ area after ablation vs. before ablation, p<0.0001, N=10. 2°-3° contact length after ablation vs. before ablation, p<0.0001, N=10. 2°-1° contact length after ablation vs. before ablation, p=0.1569, N=10. Right Panel: Fold change of the 3° cell area and cell-cell contact before and after ablation. Two-way ANOVA was performed with Sídák’s multi-comparison to compare each data set before to after ablation. 3° LCs area after ablation vs. before ablation, p<0.0001, N=8. 3°-2° contact length after ablation vs. before ablation, p=0.0015, N=8. 3°-1° contact length after ablation vs. before ablation, p<0.0001, N=8. (F) A cartoon depicting the forces affecting cell shape inferred from the pattern of recoil of 2° and 3° LCs after laser ablation. Red arrows depict contractile forces, and their thickness represents their magnitude. Green arrows depict the passive stretching of the 2°-1° cell contacts in response to the contraction of the 3° LCs. (G) After ablation of a 2° LC (marked with a yellow star), the apical area of the adjacent 3° expands. This is followed by an increase in medioapical actomyosin and cell area contraction (red arrow). Shown are snapshots from a time-lapse movie of a lattice edge before and after 2° LC ablation.

To study the role of the medioapical actomyosin network in tissue mechanics, we ablated the network either partially or entirely and investigated the cell-autonomous and non-autonomous effects. We first ablated the network in 2° LCs and found that F-actin structures recoiled toward the cell periphery (Fig. 2B, flow marked by ellipses, LC-LC contacts with white arrows). Concomitantly, we observed a rapid anisometric relaxation of the apical cell area preferentially parallel to LC-LC contacts (Fig. 2B, C, Movie 2). The severing and recoil of the medioapical actomyosin network increased the 2° LCs’ total area by approximately 2-fold, resulting from cell area expansion along the shorter axis, while the longer axis was not significantly affected (Fig. 2B-C, E). We next ablated the 3° LCs and observed a similar 2-fold increase in the apical cell area (Fig. 2D-E, Movie 2). Additionally, we found that both the LC-LC (3°-2°) and 3°-1° contacts expanded, though 3°-1° contacts expanded more (Fig. 2E). These mechanical responses suggest that the medioapical actomyosin network contracts the apical cell area asymmetrically to preferentially shrink the LC-LC and 3°-1° contacts and guide the shape changes of the LCs. Overall, these recoil patterns indicate that in 2° LCs, the medioapical actomyosin network promotes the shortening of LC-LC contacts, while in 3° LCs, it preferentially shortens the 3°-1° contact and contributes to the shortening of LC-LC contacts (see model in Fig. 2F).

The ablation of a 2° LC expanded the LC-LC contact and thus affected the adjacent non-ablated LC by stretching their shared contact. In response to this stretch, the neighboring non-ablated LC assembled a medioapical actomyosin network to contract its apical cell area and shorten the LC-LC contact (Fig. 2G). This mechanical response implies that LCs sense their mechanical state and adapt to restore proper intra- and intercellular force balance.

### Rho1 regulates the frequency and amplitude of medioapical actomyosin network assembly

We next investigated mechanisms affecting the assembly of the medioapical actomyosin network during LC remodeling. The Rho1 Rho GTPase activates effectors that promote the assembly and contractility of actomyosin networks (Ridley and Hall, 1992). Therefore, we hypothesized that the Rho GTPase cycle controls medioapical actomyosin network assembly and disassembly and, thereby, the observed fluctuations in apical cell area. To test the role of Rho1 and actomyosin dynamics in eye epithelial remodeling, we first manipulated Rho1 function itself. We reasoned that inhibiting or activating Rho1 would impair these dynamics and reveal their role in lattice remodeling. We first expressed a dominant-negative Rho1 (Rho1^DN^) and investigated the impact on medioapical F-actin dynamics. We found a general increase in apical cell areas and large fluctuations in cell area over time (Fig. 3A, yellow arrowheads mark LC-LC contacts). While both junctional and medioapical F-actin were sparse, as expected, cell area fluctuations were still observed, and contractions were accompanied by assembly of a medioapical actomyosin network that appeared to exert tension and contract overexpanded cells (Fig. 3A, red arrows). Thus, despite the impairment in F-actin levels and organization, LCs were still able to sense and adapt to their expanded state by assembling a medioapical actomyosin network to contract the apical cell area.

**Figure 3:**
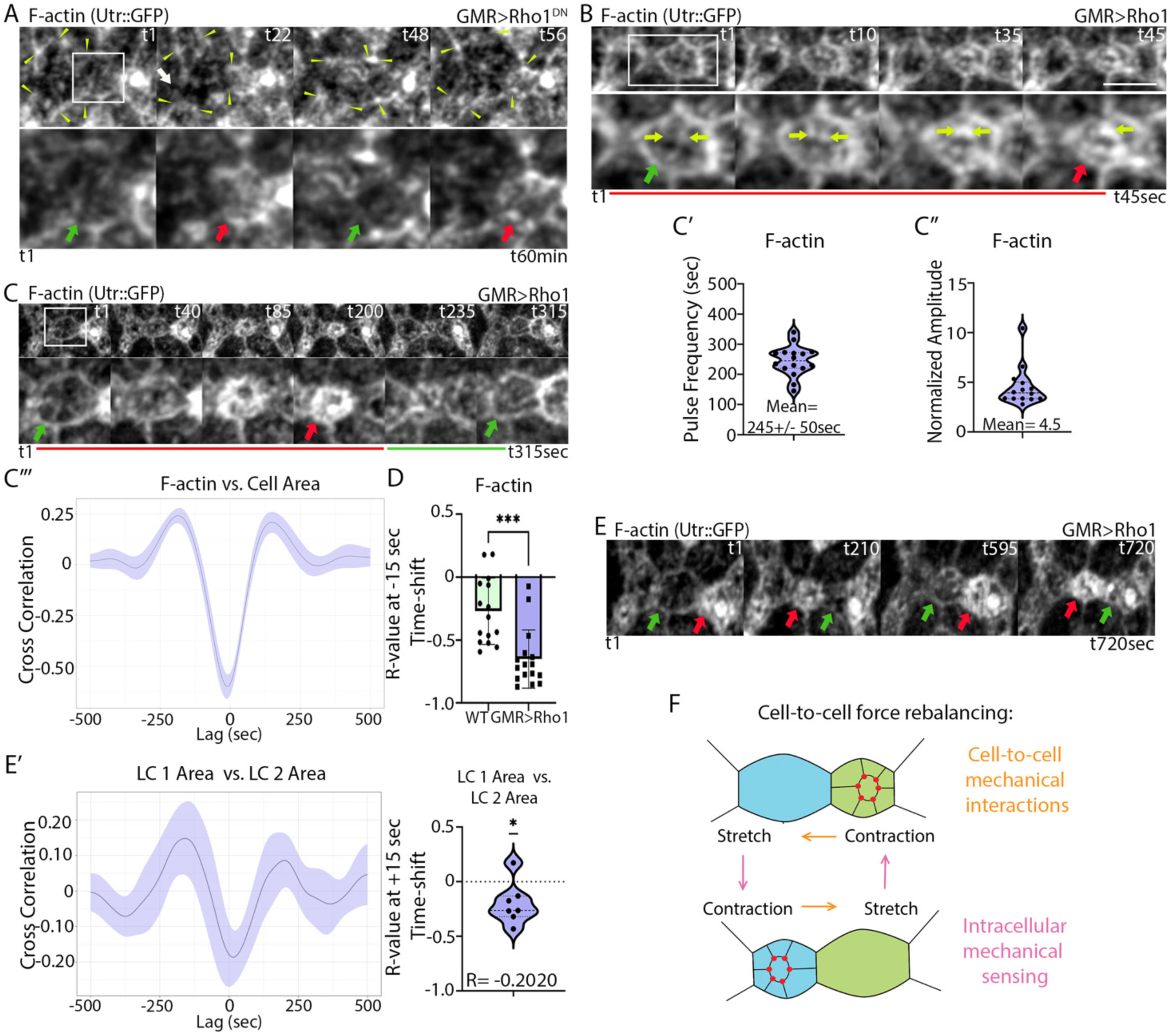
Rho1 regulates medioapical actomyosin dynamics and cell area fluctuations. (A) F-actin organization and dynamics in Rho1^DN^ expressing eyes. Top: Snapshots of a lattice edge. The continuity of the junctional actomyosin network is interrupted (white arrow). Bottom: Zoomed-in images of the 2° LCs (boxed area) show dynamic assembly and disassembly of the medioapical F-actin network correlating with cell area contraction (red arrows) and expansion (green arrows), respectively. Yellow arrowheads demarcate LC-LC contacts. (B-C) Conditional Rho1 overexpression enhances medioapical F-actin dynamics and cell area fluctuations. (B) Kymograph shows F-actin node fusion (tracked by the yellow arrows) promoting medioapical F-actin ring formation (red arrow). (C) A kymograph showing changes in medioapical F-actin in a cell that overexpresses Rho1. Green arrows, low F-actin intensity and cell area relaxation; red arrows, high F-actin intensity and cell area contraction. (C’) Medioapical F-actin pulse frequency in 2° LCs is approximately 4 minutes (245±50 sec, N=16). (C’’) F-actin levels increase on average 4.5-fold (N=14) in 2° LCs during pulsing. (C’’’) Time-shifted correlation shows that medioapical F-actin levels correlate strongly with the apical cell area contraction of 2° LCs. (D) The peak negative correlation between F-actin and cell area in Rho1 expressing eyes is significantly stronger than in WT (R=−0.6508 at peak correlation of −15sec, N=15, t-Test p=0.0002). (E) Contractile medioapical actomyosin dynamics and cell area fluctuations inversely coordinate between neighboring 2° cells prior to cell pruning. (E’) Cell area of neighboring 2° LCs is negatively correlated (R=−0.2020 at a peak correlation of +15 sec, N=14 (7 pairs), one-sample t-Test; p=0.0318). (F) A model describing the coordination of cytoskeletal and mechanical behavior between neighboring LCs.

To test whether Rho1 was sufficient to affect medioapical actomyosin dynamics, we conditionally overexpressed WT Rho1 using the GAL4/GAL80^ts^ system, successfully maintaining cell connectivity and tissue integrity. Rho1 overexpression promoted the fusion and enlargement of medioapical F-actin nodes (Fig. 3B, Movie 3). This was accompanied by increased F-actin pulse frequency (245±50 sec in Fig. 3C’, compared with 349±96 sec in WT in Fig. 1F) and amplitude (4.5-fold change, Fig. C”, compared with 1.4-fold change in WT in Fig. 1G). While in WT peak correlation between medioapical F-actin and cell area contraction was modest (Fig. 1B’’, R= −0.2706 at −15 sec), Rho1 overexpression strongly increased this correlation (Fig. 3C’’’-D, R= −0.6508 at −15 sec). These results imply that Rho1 activity is tightly regulated to control the frequency and amplitude of medioapical actomyosin assembly and contractility.

### Rho1 overexpression reveals cell-to-cell mechanical interactions

The strong fluctuations in apical cell area and medioapical actomyosin assembly induced by Rho1 overexpression prompted us to ask whether nearby cells coordinate their mechanical behavior. We focused on the behavior of adjacent 2° LCs prior to completion of cell pruning. In this cell configuration, one LC shares only one contact with the neighboring LC and a second with a mechanosensory bristle, while the second LC shares a second contact with a flanking 3° cell. Thus, adjacent LCs are exposed to the minimum of possible mechanical inputs from neighboring LCs, allowing us to isolate the mechanical interactions between these cells with low “background interference” from other neighboring cells. Inspection of time-lapse movies revealed that a contraction of a 2° LC induced a two-step response in the adjacent 2° cell (Movie 4). In the first step, the second 2° LC disassembled its medioapical actomyosin network and its apical area expanded (Fig. 3E at 210 sec, green arrow). In the second step, the second 2° LC reassembled its medioapical network and contracted its apical cell area (Fig. 3E at 595 sec, red arrow). In turn, the contraction of the second 2° LC induced the same two-step response in the first 2° LC. These cell-to-cell mechanical interactions resulted in frequent inversely coordinated cycles of expansion and contraction in adjacent LCs. The peak expansion of a LC occurred 15 sec after the maximal contraction of the adjacent LC (Fig. 3E’, R= −0.2020). Thus, as one cell contracted, the other expanded, and vice versa (Fig. 3F; Movie 4). We also observed inversely coordinated cycles of expansion and contraction between adjacent anterior and posterior cone cells and 1° cells (Fig. S2A-B’ and C-D, respectively). Thus, the medioapical actomyosin network not only contributes to cell-autonomous constriction but also exerts force on adjacent LCs that sets up anticorrelated cycles of constriction and relaxation in adjacent cells.

### Dynamic activation of MyoII and F-actin is required for tissue integrity and remodeling

Our findings suggested that tissue remodeling requires a pulsatile medioapical network controlled by Rho1 (Fig. 4A). To address this directly, we tested if MyoII and F-actin dynamics contributed to proper remodeling. To investigate the role of MyoII dynamics in this process, we expressed a constitutively active myosin light chain kinase (MLCK^CA^), which constitutively activates MyoII, and examined its effects on cellular and cytoskeletal behaviors compared with WT. Forces generated by the cytoskeleton are transmitted between cells via bicellular and tricellular adherens junctions (bAJs and tAJs, respectively). To determine the effects of constitutive MyoII activation on cell adhesion, we examined the adherens junctions (AJs) in eyes that conditionally expressed MLCK^CA^ using the GAL4/GAL80^ts^ system. We live imaged α-Catenin tagged with GFP (α-Cat::GFP) to highlight the AJs and found that they were fragmented compared with WT (Fig. 4B-C). A subset of the LCs failed to intercalate and clustered around mechanosensory bristles (Fig. 4C, yellow arrow). We also observed cells that dramatically expanded their apical area, suggesting that they failed to hold tension exerted by the medioapical actomyosin network and were pulled by their neighbors, thus compromising tissue integrity (Fig. 4C, dotted lines). MLCK^CA^ expression decreased cells’ apical area (Fig. 4E). Live imaging with MyoII::mCh showed a high level of MyoII along the entire cell perimeter. Additionally, MyoII enriched in a medioapical ring that dynamically formed and disappeared (Fig. 4F, Movie 5) on average every 10 minutes (Fig. 4F’), and its assembly correlated with cell area contraction (Fig. 4F’’, R= −0.2126). The defects that arise from MLCK^CA^ expression and constitutive MyoII phosphorylation imply that dynamic MyoII phosphorylation is required for the spatiotemporal control of actomyosin assembly, cell shape changes and epithelial integrity. This is consistent with the role of MyoII phosphorylation in other systems (Jordan and Karess, 1997; Kasza et al., 2014; Vasquez et al., 2014).

**Figure 4:**
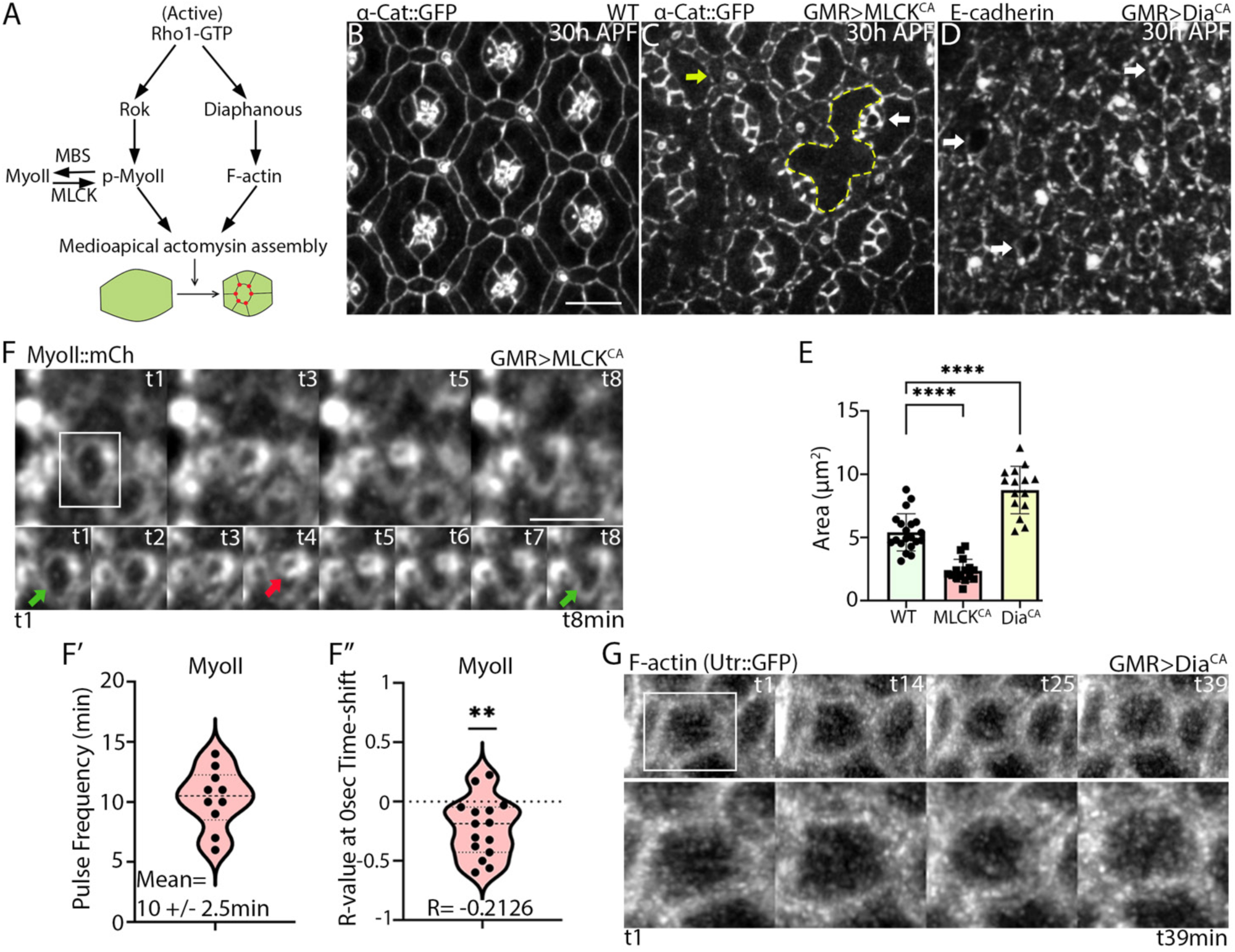
MyoII and F-actin turnover affects medioapical contractile dynamics. (A) Regulation of actomyosin contractility. (B-D) Apical cell outlines labeled with α-cat::GFP in (B) WT, (C) MLCK^CA^ and (D) Dia^CA^ expressing eyes. (C) MLCK^CA^ expression disrupted cell intercalation resulting in clustering of 2° LCs around mechanosensory bristles (yellow arrow). The AJs appeared fragmented, holes formed in the tissue (marked with yellow dotted lines) and cone cells cavitated (white arrow). (D) Dia^CA^ expression also caused fragmentation of AJs and cavitation of cone cells. (E) MLCK^CA^ expression significantly decreased the apical cell area of 2° LCs (t-test, p<0.0001, N=15), while Dia^CA^ expression increased apical cell area compared with WT (t-test, p<0.0001, N=15). (F) MyoII dynamics in a MLCK^CA^-expressing eye. A medioapical MyoII ring repeatedly contracts and expands. (F’) The frequency of medioapical MyoII ring pulsing is approximately 10 minutes±2.5 min (N=10, 30 pulses). (F’’) Medioapical MyoII levels and the apical area of 2° LCs negatively correlate (R= −0.2126, One-sample t-Test, p=0.0044, N=15) (G) F-actin dynamics and organization in Dia^CA^ expressing eyes. F-actin forms a wide band at the cell periphery and medioapical F-actin is missing.

To determine if F-actin dynamics are also required for eye epithelial remodeling, we expressed a constitutively active form of Dia (Dia^CA^), which assembles linear actin filaments downstream of Rho1, and examined the resulting phenotypes. We expressed Dia^CA^ conditionally using the GAL4/GAL80^ts^ system. First, we fixed Dia^CA^-expressing eyes and stained them for E-cad to examine the integrity of the AJs and epithelial organization. We found that the AJs were fragmented in Dia^CA^-expressing eyes like in MLCK^ca^-expressing eyes. Dia^CA^ expression also severely affected cell shape and arrangement in ommatidia (Fig. 4D). We also observed intercalation defects and cavitation of cone cells similar to those observed in eyes expressing MLCK^CA^ (Fig. 4D, white arrows). Unexpectedly, unlike the contraction of LCs in MLCK^CA^ eyes, Dia^CA^ expression significantly increased cells’ apical area (Fig. 4E). To determine if Dia^CA^ affects medioapical actomyosin, we analyzed F-actin organization and behavior in these eyes (Fig. 4G, Movie 5). In WT, F-actin is enriched both medioapically and at LC-LC contacts, while in Dia^CA^-expressing eyes, it accumulated uniformly in a thick band at the cell periphery and was missing medioapically. It is plausible that by binding efficiently to Rho1 through its RhoGTPase-binding domain, Dia^CA^ inhibits Rok activation and MyoII phosphorylation, and thus the assembly of the medioapical actomyosin network. Overall, these observations imply that medioapical MyoII and F-actin dynamics maintain a force balance in the epithelium, preserving cell shape and arrangement. This further suggests not only that Rho1 levels must be carefully controlled, but also a balance between network assembly and disassembly is required for epithelial remodeling and integrity.

### RhoGEF2 and RhoGAP71E control medioapical actomyosin dynamics

We hypothesized that a Rho1 guanine nucleotide exchange factor (RhoGEF) and Rho GTPase activating protein (RhoGAP), which respectively activate and deactivate Rho1, could regulate its medioapical dynamics (Etienne-Manneville and Hall, 2002; Jaffe and Hall, 2005). To identify these putative regulators, we carried out an RNAi screen for RhoGEFs and GAPs that influence eye epithelial remodeling. In this screen, we identified *RhoGEF2* and *RhoGAP71E* based on RNAi phenotypes in epithelial remodeling. This led us to hypothesize that both activation of Rho1 by RhoGEF2 and inhibition by RhoGAP71E regulate Rho1’s GTPase cycle and drive medioapical pulsatile dynamics during remodeling. We tested these ideas below.

### RhoGEF2 accelerates the frequency and amplitude of medioapical actomyosin assembly and fluctuations in apical cell area

To determine the site of RhoGEF2 action, we examined RhoGEF2 localization using a RhoGEF2 reporter tagged with GFP (RhoGEF2::GFP) and marked cell outlines using F-actin with Lifeact::Ruby (Mason et al., 2016). RhoGEF2::GFP accumulated medioapically in pulses (Fig. 5A, yellow arrows) that partially overlapped with medioapical F-actin. To determine if RhoGEF2 levels correlate with apical cell area, we measured the correlation between RhoGEF2 levels and cell area over time and found a weak negative correlation (Fig. 5A’, left panel), with peak enrichment of RhoGEF2 occurring 15-20 seconds after peak contraction (Fig. 5A’, right panel, R= −0.1361).

**Figure 5:**
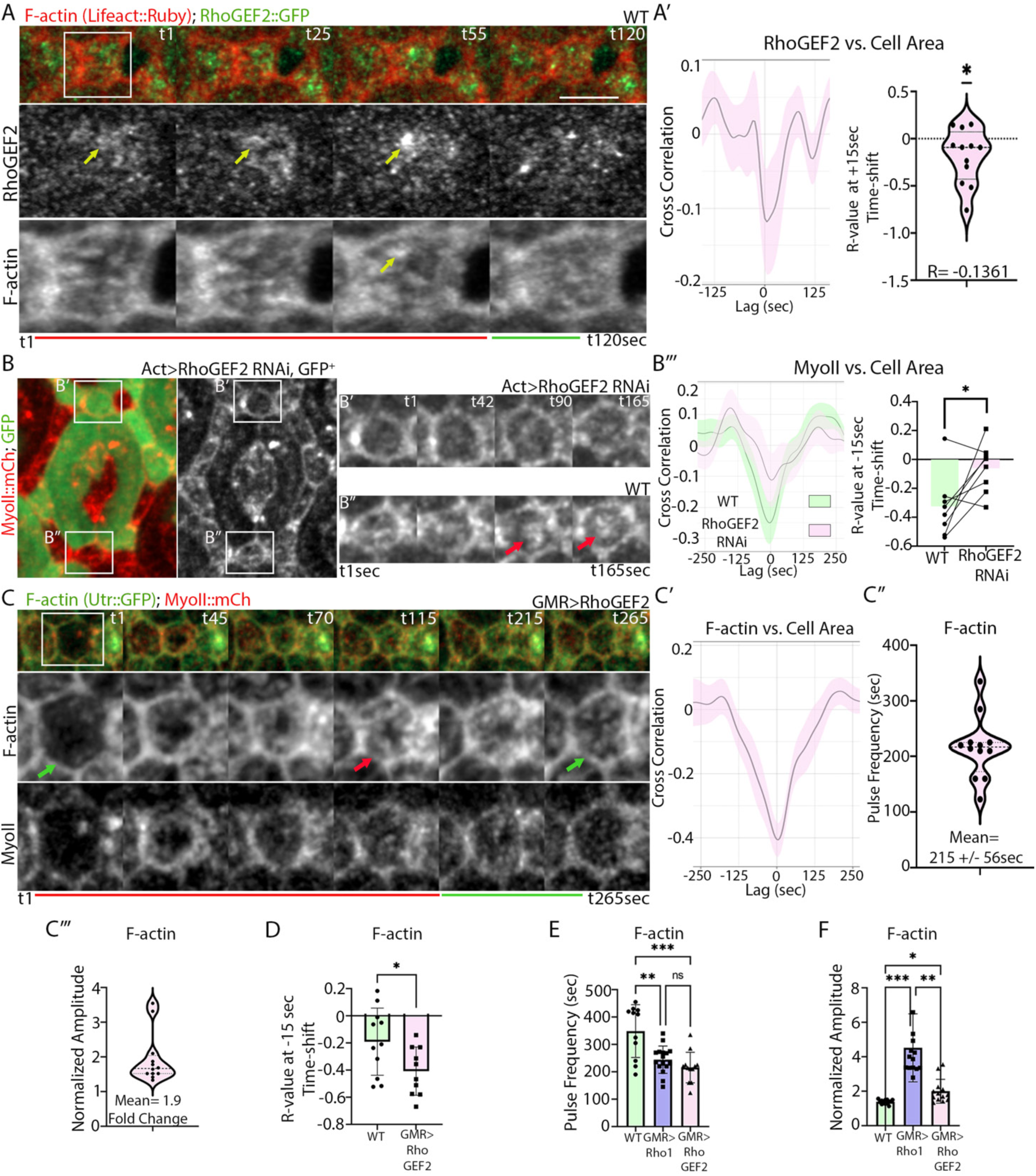
RhoGEF2 promotes the assembly of the medioapical actomyosin network. (A) RhoGEF2::GFP and F-actin labeled with Lifeact::Ruby. Snapshots from a time-lapse movie of a lattice edge (top) and magnified views of a 2° LC demarcated with a box. RhoGEF2::GFP (yellow arrow) accumulates medioapically in pulses. (A’) Left: Time-shifted correlation plot between RhoGEF2::GFP and apical cell area. Right: RhoGEF2::GFP accumulation correlates negatively with cell area at a peak of +15 sec (R=−0.1361, N=12, one-sample t-Test p=0.0250). (B-B’’’) Clonal depletion of *RhoGEF2* by RNAi reduces medioapical MyoII assembly compared with the WT counterparts. (B’) Kymograph showing a reduction in MyoII levels in the RhoGEF2RNAi-expressing cells compared with WT (B’’). (B’’’) Left: Time-shifted correlation plots between medioapical MyoII levels and LC area in RhoGEF2RNAi-expressing cells (pink) compared with WT (green). Right: Peak correlation is weaker in RhoGEF2RNAi-expressing cells compared with WT counterparts (paired t-test, p= 0.0281, N=8 pairs). (C) RhoGEF2 overexpression enhanced medioapical actomyosin network assembly and apical cell area fluctuations. Top: Snapshots of a lattice edge. Bottom: Magnified views of the 2° LC show the assembly (red arrow) and disassembly (green arrow) of the medioapical actomyosin network that correlate with cell area contraction and expansion, respectively. (C’) Medioapical F-actin levels correlate with apical area contraction in 2° LCs. RhoGEF2 overexpression increases (C’’) medioapical F-actin pulse frequency to approximately 3 minutes (215±56 sec, N=12), and (C’’’) normalized amplitude. (D) RhoGEF2 expression increases the peak correlation between F-actin and cell area (peak correlation at −15 sec, N=11, t-Test p=0.0337). (E) Rho1 and RhoGEF2 expression increase the pulse frequency and (F) pulse amplitude of F-actin accumulation compared with WT. (E) Pulse frequencies of medioapical F-actin in WT eyes compared to eyes overexpressing Rho1 and RhoGEF2. One-way ANOVA with Tukey’s multiple comparisons. WT vs. GMR>Rho1, difference of means=104 sec, p=0.0010. WT vs. GMR>RhoGEF2, difference of means=133 sec, p=0.0001. GMR>Rho1 vs. GMR>RhoGEF2, difference of means=29 sec, p=0.5084, N=11-16. (F) Pulse amplitudes of F-actin in the above genotypes. Welch One-way ANOVA with Dunnett’s comparison between WT and GMR>Rho1, mean difference of R=−0.3139, p<0.0001. WT vs. GMR>RhoGEF2, mean difference of R=−0.6155, p=0.0218. GMR>Rho1 vs. GMR>RhoGEF2, mean difference of R=2.523, p= 0.0011, N=14.

To test if RhoGEF2 controls medioapical actomyosin dynamics, we examined F-actin and MyoII dynamics in RhoGEF2RNAi-expressing clones (Fig. 5B-B’’’, Movie 6). We found a decrease in medioapical actomyosin levels in the clones (Fig. 5B’, Movie 6) compared with WT counterparts (Fig. 5B’’). There was a weaker peak correlation (at −15 sec) between MyoII accumulation and apical cell area contraction in RhoGEF2 RNAi clones compared with WT counterparts (Fig. 5B”’). These results provide evidence that RhoGEF2 activates Rho1 to control the assembly of the medioapical actomyosin network.

We also investigated the impact of RhoGEF2 overexpression on these dynamics. We found that RhoGEF2 overexpression induced robust and more frequent fluctuations of the medioapical actomyosin network that coincided with changes in the cell apical area (Fig. 5C, Movie 6). Actomyosin assembled into a dense network that correlated with apical area contraction (red arrow) and then disassembled (green arrow), resulting in apical area relaxation. The assembly of the medioapical actomyosin network strongly correlated with the apical cell area contraction (Fig. 5C’, R= −0.3688). While in WT, medioapical F-actin correlates with apical cell area contraction (Fig. 1B’’, Fig. S1B’’, R= −0.2706), RhoGEF2 overexpression enhanced this correlation (Fig. 5C’, D, R= −0.4076). Furthermore, overexpression of RhoGEF2 accelerated F-actin pulse frequency (Fig. 5C’’, 215±56 sec) compared with WT (Fig. 5E, 349±96 sec), and enhanced the amplitude of F-actin accumulation (Fig. 5C’’’) compared with WT (Fig. 5F). Thus, RhoGEF2 overexpression increased the amplitude and frequency of medioapical actomyosin assembly and phenocopied the Rho1 overexpression phenotypes. The results suggest that RhoGEF2 activates Rho1 to promote the assembly of a contractile medioapical actomyosin network during LCs remodeling.

### RhoGAP71E promotes the disassembly of the medioapical actomyosin network

To determine where in the cell RhoGAP71E acts, we live-imaged a RhoGAP71E reporter tagged with GFP (RhoGAP71E::GFP) (Denk-Lobnig et al., 2021). RhoGAP71E::GFP was enriched at the cell surface and accumulated medioapically during cell area contractions (Fig. 6A, yellow arrow). RhoGAP71E accumulated maximally 10 seconds after peak LC contraction and levels decreased during apical area expansion (Fig. 6A’, R= −0.1399).

**Figure 6:**
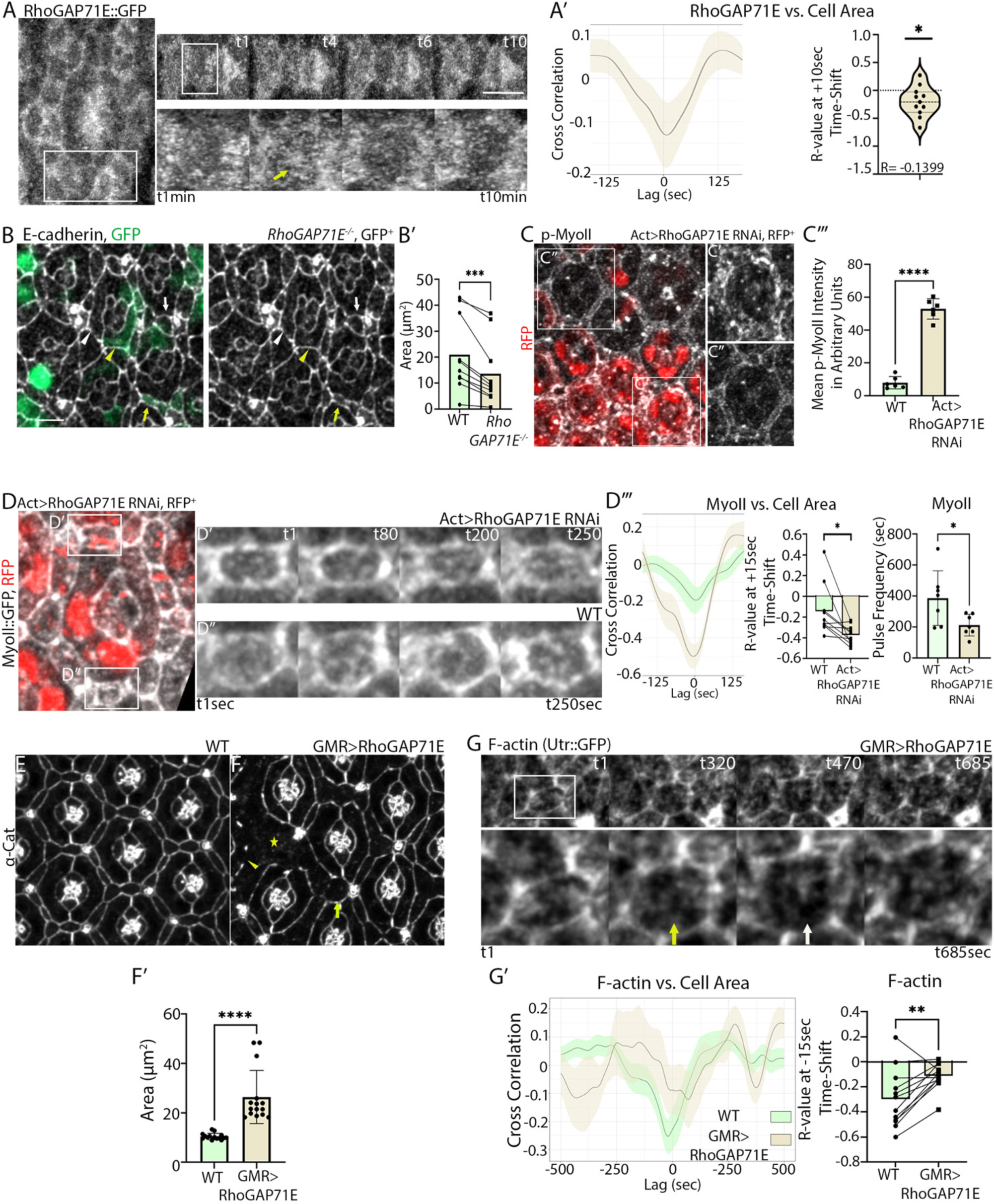
RhoGAP71E promotes the disassembly of the medioapical actomyosin network. (A) RhoGAP71E::GFP accumulated medioapically in pulses (yellow arrow). Snapshots from a time-lapse movie of an ommatidium (left), lattice edge (top), and magnified views of a 2° LC demarcated with a box. (A’) Medioapical RhoGAP71E::GFP levels negatively correlate with apical cell area of 2° LCs at a time shift of +10 sec (R= −0.1399, N=11, one-sample t-Test p=0.0328). (B-D) RhoGAP71E promotes apical cell area expansion. (B) *RhoGAP71E* mutant cells positively marked by GFP in fixed tissue stained for E-cadherin. White arrow, WT LCs; yellow arrow, mutant LC; white arrowhead, WT 1° cells; yellow arrowhead, mutant 1° cell. (B’) The apical area of the *RhoGAP71E* mutant cells is significantly smaller than the area of WT counterparts (Paired t-Test, p=0.0007, N=10 pairs). (C) RhoGAP71E RNAi expression decreases MyoII phosphorylation. (C’) p-MyoII levels are higher in the clones (C’’) compared with the WT regions. (C’’’) T-test, p<0.0001, N=6. (D) RhoGAP71E RNAi expression decreases apical cell area and the correlation between MyoII accumulation and cell area contraction. Left: Snapshot from a time-lapse movie of MyoII (Sqh::GFP) in eyes with RhoGAP71E RNAi-expressing clones (RFP^+^). Kymographs of zoomed-in views of (D’) a RhoGAP71E RNAi-expressing 2° LC compared with (D’’) a WT counterpart. (D’’’) Peak correlation (at −15 sec) between MyoII accumulation and apical area contraction is higher in RhoGAP71E RNAi-expressing cells (light brown) compared with WT (green). Paired t-test, p=0.0127, N=10 pairs. Right: RhoGAP71E RNAi expression increased medioapical MyoII pulse frequency in the 2° LC to approximately 3 minutes (212±69 sec) compared with approximately 6.5 minutes in WT (387±177 sec, t-test, p= 0.0109, N=7). (F) RhoGAP71E overexpression disrupted epithelial organization (yellow star) and caused expansion of the cells’ apical area (yellow arrow) and gaps in the continuity of the AJs (yellow arrowhead), compared with (E) WT. (F’) RhoGAP71E expression significantly increased the apical cell area of 2° LCs compared with WT (N=14, t-Test, p <0.0001). (G) RhoGAP71E overexpression reduced medioapical F-actin levels. Top: Snapshots of a lattice edge. Bottom: A magnified view of the boxed 2° LC. Yellow arrow, a faint actomyosin ring can be detected in LCs that failed to contract their apical cell area. White arrow, network disassembly did not lead to cell area expansion. (G’) Left: RhoGAP71E overexpression abolished the correlation between actomyosin accumulation and cell area contraction. Right: Paired t-test at a time shift of −15 sec (p=0.0060, N=12 pairs).

To determine if RhoGAP71E controls medioapical actomyosin dynamics, we first examined the effects of *RhoGAP71E* loss on LCs’ remodeling. Eliminating *RhoGAP71E* function or expressing RhoGAP71E RNAi in whole eyes produced flies without eyes. However, using the FLP/FRT technique, we were able to recover sparse *RhoGAP71E* mutant cells (Fig. 6B). The *RhoGAP71E* 1° and 2° mutant cells (yellow arrowhead and arrow, respectively) had smaller apical areas compared with WT counterparts (white arrow and arrowhead, respectively, Fig. 6B’). This observation suggested that actomyosin assembly and tension exerted on the apical cell perimeter were higher in the *RhoGAP71E* mutant cells, leading to apical area contraction. To confirm this result, we examined levels of the phosphorylated activated form of MyoII (p-MyoII) in clones expressing RhoGAP71E RNAi (Fig. 6C). We found an increase in p-MyoII levels in the clones (Fig. 6C’) compared with WT cells (Fig. 6C’’ and C’’’). We also live imaged clones expressing RhoGAP71E RNAi (Fig. 6D, Movie 7). Opposite to the effect of RhoGEF2 depletion on cell shape, we found a decrease in the apical cell area in cell clones expressing RhoGAP71E RNAi (Fig 6D’) compared with WT cells (Fig. 6D’’). The increased assembly of the medioapical actomyosin network in the RhoGAP71E RNAi-expressing cells increased the correlation between MyoII and cell area contraction (Fig. 6D’’’), and also led to increased MyoII pulse frequency compared with WT (Fig. 6D’’’, 212±69 sec and 387±177 sec, respectively). These results suggest that RhoGAP71E inhibits medioapical actomyosin network assembly during LCs remodeling, as decreasing RhoGPA71E levels increased the amplitude and frequency of medioapical MyoII assembly.

We also examined the effects of RhoGAP71E overexpression on medioapical actomyosin and apical cell area fluctuations. RhoGAP71E overexpression expanded the apical cell area and compromised the ability of a subset of cells to hold tension resulting in their expansion (Fig. 6E-F’, yellow arrow). We also observed a loss of continuity of AJs marked with α-Cat::GFP in expanded cells (Fig. 6F, yellow arrowhead). These results imply that RhoGAP71E overexpression reduced medioapical tension, allowing the apical cell perimeter to relax. Although medioapical F-actin fluctuated in these eyes (Fig. 6G, yellow and white arrows, Movie 7), it appeared to be less robust, and network assembly did not correlate with cell area contraction as in WT eyes (Fig. 6G’). Finally, we examined the interaction between RhoGAP71E and RhoGEF2 by co-expressing the two proteins in the retina. While RhoGAP71E expression alone expanded the apical cell area and led to cell intercalation defects, the expression of RhoGEF2 with RhoGAP71E suppressed these phenotypes (Fig. S3A-D). Together, these findings provide evidence that RhoGAP71E modulates medioapical actomyosin dynamics and tension by inhibiting Rho1 activity medioapically and antagonizing RhoGEF2 function. Thus, RhoGAP71 and RhoGEF2 regulate the Rho1 GTPase cycle and the overall levels of Rho1 activity to control medioapical actomyosin assembly and contractility, which govern cell shape and arrangements.

## DISCUSSION

During epithelial remodeling of the fly retina, at least two dynamic actomyosin networks operate at the apical region of the cells. The first is a junctional network that cyclically contracts and expands the LC-LC contacts (Del Signore et al., 2018; Malin et al., 2022). Here we describe a second mechanism essential for accurate morphogenesis: a non-ratcheting pulsatile medioapical actomyosin network controlled by Rho1 and its effectors Dia and MyoII. This network is established by flows of F-actin from the cell surface toward the medioapical region where it assembles into actomyosin nodes that generate anisotropic contractile force with the highest force predicting the cell edges that will eventually shorten. This force is transient, and upon release, the actomyosin nodes remodel, disassemble or merge with the junctional network. Strikingly, pulses of medioapical contraction and release are reciprocally synchronized between adjacent 2° LCs, revealing that these Rho1-regulated networks are integral to biomechanical feedback coupling. That is, they appear to trigger, and be triggered by, forces between neighboring cells and coordinate their activity. Furthermore, we found that Rho1 activity is positively regulated by RhoGEF2 and negatively by RhoGAP71E. These observations expand our understanding of these players’ roles in embryonic undifferentiated tissue development to a fully differentiated tissue where they work over different time scales and levels of cellular shape determination.

While a previous study reported the existence of medioapical actomyosin networks that can generate force in 1° cells, our results add additional insight into the properties of these networks and their role in lattice cells (Blackie et al., 2020). We show that medioapical actomyosin networks depend on Rho1 and Rho1 effectors Dia and MyoII. Perturbation of these networks not only led to remodeling errors but also to tissue rupture. These findings indicate a crucial role for medioapical actomyosin networks in the mechanical system that promotes epithelial integrity. In contrast to retina 1° cells, we find that forces generated by the actomyosin network in 2° cells are anisometric. When ablated, the largest recoil occurs parallel to LC-LC contacts, predicting the edges that will eventually shorten. This is consistent with our previous observation that single *myoII* mutant LCs change shape anisometrically (Del Signore et al., 2018).

Live imaging of LCs also revealed the assembly of a medioapical actomyosin ring structure and its dynamics. In other cases, the medioapical actomyosin network forms a ‘meshwork’ (Chanet et al., 2017; Christodoulou and Skourides, 2015; Collinet et al., 2015; He et al., 2010; Maitre et al., 2015; Martin et al., 2009; Rauzi et al., 2010; Solon et al., 2009; Yu and Fernandez-Gonzalez, 2016). This is the case in gastrulation, where cells’ resistance to cell contraction is low in one direction and high in the orthogonal direction (Martin et al., 2010). In experimentally generated round embryos, when the resistant forces are increasingly isometric, the medioapical actomyosin meshwork converts to rings (Chanet et al., 2017). Therefore, it has been postulated that isometric resistance fosters formation of a medioapical actomyosin ring while directional resistance fosters an apparent meshwork. Taken together with the current observations, a ring structure may be associated with conditions where opposing forces are generally isometric; however, this organization is, in turn, capable of generating anisometric force.

Our results also revealed the pulsing features of medioapical actomyosin networks in the retina. We found that Rho1 overexpression increased the amplitude of F-actin accumulation, area constriction, and frequency of apical constriction. This observation suggests that Rho1 activity level affects the kinetics of pulsing. However, we found that these characteristics of pulsing had no obvious impact on the persistence of constriction or the appearance of ratcheting. This is different from other systems regulated by medioapical actomyosin pulsing. For example, in ventral furrow formation, there are correlations between myosin accumulation and the frequency and strength of constriction, but these are, in turn, correlated with the likelihood of ratcheting (Chanet et al., 2017). Thus, our results show unique features of medioapical actomyosin pulsing that depend on the specific developmental process and eventual shape-change required.

Overexpressing Rho1 also dramatically revealed that the induced rapid cell area oscillations were inversely coordinated in neighboring 2° LCs: one cell contracted while its neighbor relaxed and vice versa. Although it is known that cells can sense and adapt to mechanical forces (del Rio et al., 2009; Gaertner et al., 2022; Mueller et al., 2017; Spadaro et al., 2017; Yonemura et al., 2010), and medioapical actomyosin generates tension in the retina (Blackie et al., 2020), the current observations expand our understanding of how forces work in this system. The data suggest that neighboring LCs and the medioapical actomyosin networks do not merely strike a balance of mechanical forces but are continually adapting to changing forces over time, generating coordinated pulsing changes in cell shape. More specifically, our findings provide evidence that mechanical cell-to-cell interactions can activate or inhibit Rho1 function and actomyosin contractility and that cells respond to pull and stretch under normal physiological conditions by modulating Rho1 function. Thus, Rho1-dependent medioapical actomyosin networks appear integral to feedback loops that coordinate pulsing in neighboring cells.

Not only was anti-correlated pulsing between neighboring cells detected with Rho1 overexpression, but it was also apparent in WT eyes (Movie 1). However, the correlation between actomyosin accumulation and cell area contraction in WT cells had a wide distribution (Fig. S1B”, C’). This suggests that the pulsing is highly tuned to work in balance with other ongoing cellular processes that may provide permissive conditions, such as the net force exerted by neighboring cells. The anti-correlated pulsing of neighboring 2° LCs is a unique feature contrasting with other examples of pulsatile contraction. For example, in cell intercalation during germband extension, adjacent cells constrict simultaneously, resulting in loss of contacts and rearrangement of the epithelium (Fernandez-Gonzalez and Zallen, 2011; Rauzi et al., 2010; Sawyer et al., 2011; Vanderleest et al., 2018). In apical constriction, it has been observed that when one cell exhibited ratcheting contraction, ratcheting of nearby cells became more likely, to stabilize changes in cell shape and facilitate invagination (Xie and Martin, 2015). These findings suggest that although Rho1-regulated medioapical actomyosin constriction generally appears sensitive to forces exerted across the epithelium, the cell’s specific response to forces imposed by neighbors is a control point with a variable outcome.

Together, these findings indicate that Rho1 dynamics must be tightly controlled in space and time to regulate actomyosin dynamics and epithelial morphogenesis. Using a genetic screen, we identified RhoGEF2 and RhoGAP71E as upstream regulators of Rho1 whose perturbation yielded altered cell shape while the integrity of the adherens junctions was maintained. Although we previously identified these genes in a screen for epithelial folding in leg joint morphogenesis, and they are both known regulators of Rho1 in other systems, we were surprised to identify these genes because the characteristics of the medioapical actomyosin network in the LCs are unique (Azevedo et al., 2011; Denk-Lobnig et al., 2021; Fox and Peifer, 2007; Greenberg and Hatini, 2011; Hacker and Perrimon, 1998; Mason et al., 2016; Mulinari et al., 2008). In these other cases involving RhoGEF2 and RhoGAP71E, there is typically ratcheting shape change, irreversible loss of contacts, and/or epithelial folding, none of which characterize changes in the retina at this phase of development. Thus, identifying RhoGEF2 and RhoGAP71E in this study is important because it reveals that their participation is not linked to a particular morphological outcome. Since pulsing appears regulated by forces imposed by neighboring cells, our results also suggest the hypothesis that mechanical forces could, in turn, regulate RhoGEF2 and/or RhoGAP71E. In sum, these results provide a starting point for the revelation of the regulatory networks that guide these intermediaries to differential developmental outcomes.

Furthermore, our data indicate that tuning of pulsing dynamics by Rho1 and its regulators is crucial for the cells to maintain their shape and position in the lattice and prevent catastrophic tissue ruptures during this period of eye development. However, it is interesting to note that with constitutively active MLCK, we continued to observe pulsing accumulation of MyoII, even though the tissue architecture was highly disrupted. This observation is consistent with the idea that a core of pulsing dynamics may arise from the intrinsic biophysical properties of contractile actomyosin networks, wherein as F-actin filaments are exposed to high tension, they spontaneously disassemble (Haviv et al., 2008; Munjal et al., 2015). However, our results would also support the idea that pulsing characteristics are finely tuned to work in balance with other processes occurring in the epithelium to maintain tissue structure and generate the appropriate developmental pattern.

A salient feature of retinal development is the continual movement and shape changes of cells, even though it takes hours for persistent changes to unfold. Our results suggest that continuous cycling of the nucleotide state of the Rho1 GTPases drives Rho1-dependent regulation of the cytoskeleton that underlies components of these movements. Thus, these are active, energy-dependent processes that cells invest in, despite the apparent transience of shape changes seen over the short term. There are various explanations for why medioapical actomyosin pulsing exists in other systems, for example, to enable apical constriction by ratcheting or to utilize the relaxation phase for plasma membrane remodeling to remove membrane and junctional proteins (Jewett et al., 2017; Miao et al., 2019). Here in the *Drosophila* retina, it appears medioapical actomyosin pulsing coordinates the mechanical behavior of neighboring cells and functions to maintain mechanical tissue integrity. Previously, we described oscillations of junctional networks of LC-LC contacts that occurred with a periodicity of about 15 minutes (Del Signore et al., 2018; Malin et al., 2022). Here, pulsing of medioapical actomyosin networks has a periodicity closer to 6 minutes. While we do not yet know how these processes interplay, it is intriguing to note the existence of multiple sinusoidal functions that could distribute mechanical forces and speculate on how their actions are eventually integrated to generate the precise shaping of the eye.

In summary, we identified a pulsatile medioapical actomyosin network in LCs that exerts tension on the apical cell surface in a pattern that predicts the final shape changes of the cells. This network assembles into nodes connected by F-actin filaments that exert tension on one another and the cell surface. The network assembly is controlled by cell-to-cell mechanical interactions and is modulated intracellularly by RhoGAP71E and RhoGEF2. These mechanical cell-to-cell interactions manifest in inversely synchronized cell-to-cell oscillations of actomyosin and the contraction of apical cell area (Fig. 7). Our research suggests that during epithelial remodeling, when the cells are engaged in the pursuit of stable forms and interaction with neighbors, finely tuned pulsing of medioapical actomyosin networks functions in balancing the forces between epithelial cells. Regulated by mechanical and biochemical signals, Rho1 plays a crucial role in this delicate rebalancing act.

**Figure 7:**
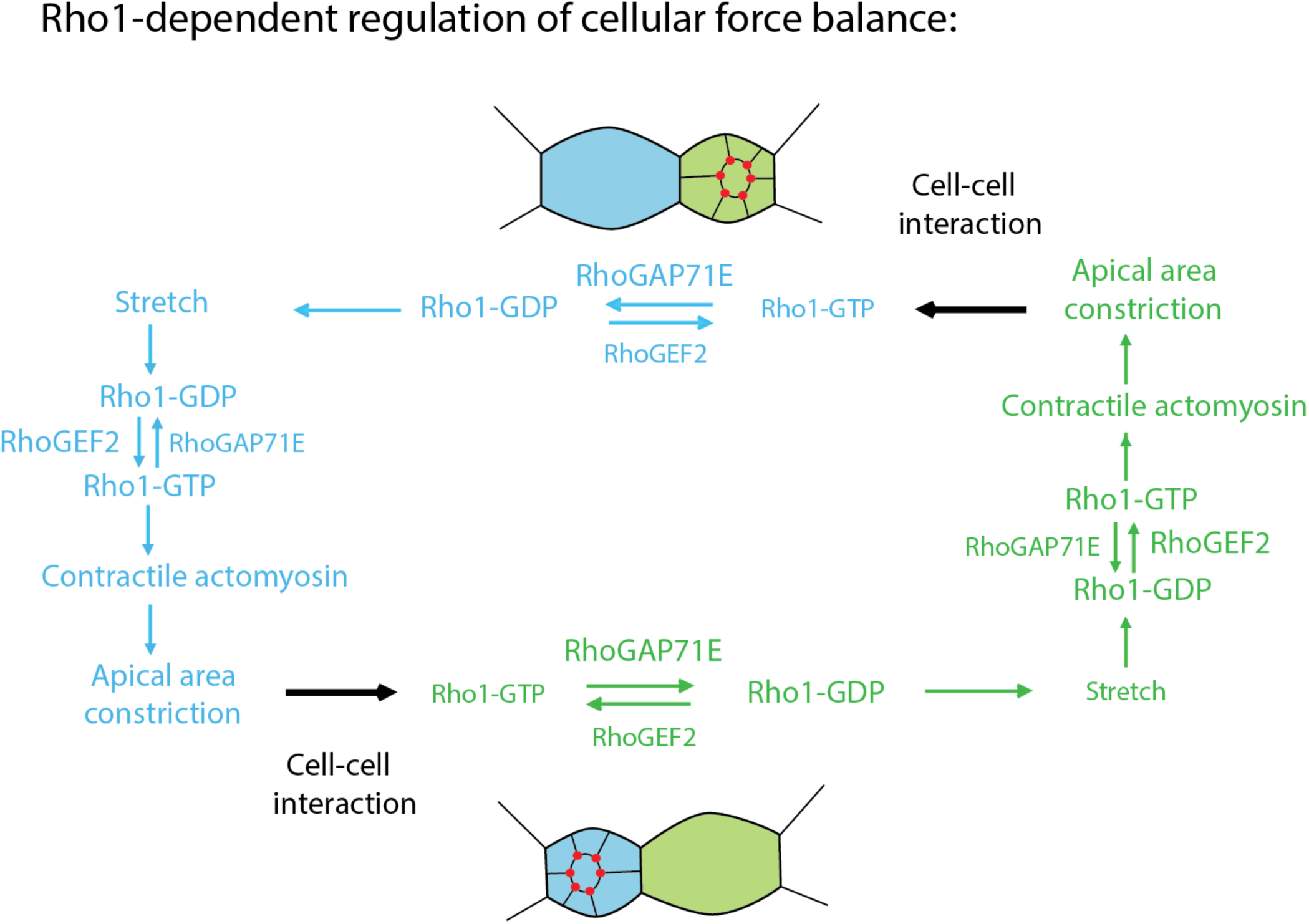
Regulated cell-to-cell coordination of mechanical behavior. A summary diagram of the results reported in this study. LCs are mechanically coupled, and their mechanical behavior is inversely coordinated. The contraction of a LCs stretches the adjacent LC and promotes the disassembly of its contractile medioapical actomyosin network. This is sensed by the stretched cell and leads to the assembly of a contractile network that contracts its apical area. Cell area contraction stretches the neighboring cells’ apical area, promoting the disassembly of its medioapical actomyosin network, followed by reassembly to repeat the cycle. This cell-to-cell coordination of cytoskeletal and mechanical behavior is controlled by and is dependent on dynamic Rho1 function and downstream effectors Dia and MyoII. Rho1 must be both activated by RhoGEF2 and inhibited by RhoGAP71E for these dynamics to occur.

## ACKNOWLEDGEMENTS

We thank J. Treisman, U. Tepass, A. Martin and J. Zallen for generous gifts of flies, the Bloomington *Drosophila* Stock Center, the Vienna *Drosophila* Research Center and the Kyoto Stock Center for flies, the Developmental Studies Hybridoma Bank, R. Ward and J. Treisman for antibodies, and G. Rong and the Institute for Chemical Imaging of Living System (CILS) at Northeastern University for assistance with multi-photon imaging. We thank K. G. Commons for her critical reading of the manuscript and editorial suggestions and Paul Hatini for writing the R script to perform the time-shifted Pearson’s correlation analysis and plot the results.

This work was supported by a grant from the National Institute of Health to V.H. (R01 GM129151).

The authors declare no competing financial interests.

## Author Contribution

Conceptualization, V. Hatini; Investigation and Resources, C. Rosa, J. Malin and V. Hatini; Formal Analysis, C. Rosa; Writing Original Draft, V. Hatini; Manuscript Review and Editing, C. Rosa, J. Malin and V. Hatini; Supervision, V. Hatini; Funding Acquisition, V. Hatini.

## Data availability

All data needed to evaluate the conclusions in the paper are present in the article and/or the supplementary materials. All image data will be deposited in the Tufts Metaverse database and given an accession number prior to publication. Image data and other supporting data of this study are available from the corresponding author upon reasonable request. Requests for reagents should be directed to and will be fulfilled by the lead contact, Victor Hatini.

## METHODS

### Fly strains

We employed a RhoGEF2::GFP BAC transgene inserted at the VK33 attP transgene landing site to examine RhoGEF2 dynamics during LCs remodeling (Mason et al., 2016). To examine the RhoGEF2 overexpression phenotypes, we expressed UAS-RhoGEF2 protein with an N-terminal T7 tag. We employed a RhoGAP71E::GFP CRISPR insertion (Denk-Lobnig et al., 2021) and RhoGAP71E::GFP driven by the ubiquitin promoter (this study) to examine RhoGAP71E protein distribution and dynamics, a WT UAS-RhoGAP71E (this study) to examine overexpression phenotypes and *RhoGAP71E*^j6B9^ FRT80B to generate genetically marked mutant clones.

Fly lines from the Bloomington *Drosophila* Stock Center: (1) Rho1::GFP, (2) UAS-Rho1.N17, (3) UAS-Rho1 (4) UAS-MLCK.Ct, (5) UAS-Dia.CA, (6) UAS-T7.RhoGEF2, (7) UAS-Lifeact::Ruby, (8) sqh-GFP::Rok, (9) GMR-GAL4, (10) *y w*; Actin>y+>GAL4, UAS-GFP, (11) *w;* Actin>CD2stop>GAL4, UAS-mRFP, (12) *y w*; α-Cat::GFP^gfstf^, (13) *y w* hsFLP; UAS-GFP^nls^, (14) *w;* tub-GAL4; FRT40A, tub-GAL80^ts^, (15) *w;* FRT80B, Ubi-GFP, (17) UAS-RhoGEF2-RNAi (BL-34643) and (18) UAS-RhoGAP71E-RNAi (BL-32417). α-Cat::Venus^CPTI002596^ and *RhoGAP71E ^j6B9^* FRT80B were obtained from the Kyoto Stock Center. Fly lines previously described by Del Signore et. al. (2018) and Malin et al. (2022): (1) UAS-Lifeact::Ruby; GMR-GAL4, (2) UAS-Lifeact::GFP; Sqh-Sqh::mCherry, GMR-GAL4, (3) UAS-Lifeact::Ruby; Sqh-Rok::GFP, GMR-GAL4. Additional stocks used: (1) Sqh-Sqh::mCherry and (2) Sqh-UtrABD::GFP, Sqh-Sqh::mCherry (gift of A. Martin), (3) GMR-GAL4, UAS−α-Cat::GFP (gift of R. Cagan), (4) RhoGAP71E::GFP and (5) RhoGEF2::GFP (gift of A. Martin).

### Molecular biology and construction of genetically encoded reporters

We generated several new reporters driven by the UAS and the ubiquitin promoter including Ubi-RhoGAP71E::GFP and UAS-RhoGAP71E::GFP. The *RhoGAP71E* open reading frame (ORF) was amplified from cDNA clone LD04071. The PCR products were inserted into the pENTR plasmid by Topo cloning. All expression clones were generated by the Gateway technology using the *Drosophila* Gateway Vector Collections (gift of T. Murphy and C.-Y. Pay) using the Clonase II reaction to fuse the ORFs in frame with a desired fluorescent protein. Transgenic flies carrying these transgenes were established by standard methods by BestGene Inc.

### Genetic analysis

GMR-GAL4 was used to broadly express UAS-transgenes in the eye (Wernet et al., 2003). The FLP/FRT (Xu & Rubin, 1995) and MARCM techniques (Lee and Luo, 2001) were used to generate genetically marked clones by FLP-mediated mitotic recombination and the FLP-Out/GAL4 technique to express desired transgenes in genetically marked clones. hsFLP; Ubi-GFP, FRT80B was used to generate *RhoGAP71E* mutant FLP/FRT clones. Mitotic and FLP-Out clones were induced by a heat shock for 30 minutes at 34°C.

## METHOD DETAILS

### Immunofluorescence

White prepupae (0h APF) were selected and aged on glass slides in a humidifying chamber at 25°C. Pupal eyes were dissected in phosphate-buffered saline, fixed for 35 minutes in 4% paraformaldehyde in PBS and stained with antibodies in PBS with 3% BSA, 0.3% Triton X-100 and 0.01% Sodium Azide. Primary antibodies used were rat anti-E-cad (DSHB #DCAD2, 1:100), mouse anti-Dlg (DSHB #4F3, 1:500) and guinea pig anti-Sqh1P (gift from R. Ward). Alexa 405 (Calbiochem), Alexa488, Alexa647 (Molecular Probes), and Cy3 and Cy5 (Jackson ImmunoResearch) conjugated secondary antibodies were used at 1:150.

### Confocal time-lapse imaging

Flies were selected for imaging at 0h APF and aged at 25°C unless described otherwise. Pupae were mounted in a slit created in an agarose block with eyes facing the coverslip after the operculum for each pupa was removed. The agarose block was surrounded by a Sylgard 184 gasket prepared in the lab and capped with a custom-built humidified chamber. Time-lapse imaging was performed on a Zeiss LSM800 inverted laser scanning confocal microscope with Airyscan. Images were taken every 5, 30, and 60 sec as noted with an optimal pinhole (1 AU) using a 63x, 1.4-NA, plan Apochromat immersion objective, 0.7 μm per optical section with a 10–50% overlap between sections, at a scan speed of 6-7 with no averaging.

### Laser ablation

Laser nano-ablations to probe tissue mechanics were performed using a Zeiss LSM880 NLO with a near-infrared InSight X3 two-photon tunable laser (680-1300nm) using a 740nm multiphoton excitation. Samples were imaged with Utr::GFP and Sqh::mCh to identify the medioapical actomyosin network for ablation using a 63X oil immersion objective (N.A 1.4). A small field size, of 4×4 pixels, was ablated in the center of either 2° or 3° LCs using a 30-40% laser output, at a scan speed of one, with one iteration. Images were collected every second for 250 sec in a frame of 1584×1584 pixels. Ablations were performed 6 seconds after the beginning of each experiment.

### Quantification and Statistical Analyses

#### Cell expansion analysis

The areas of 2° and 3° LCs were measured in Fiji immediately before and 180 seconds after ablation to calculate the fold expansion of the apical cell area. Ratios were calculated in Microsoft Excel by dividing the area after ablation by the area before ablation. LC-LC (2°-3°), 2°-1°, and 3°-1 contact lengths for both 2° and 3° LCs were also measured before ablation and 180 seconds after ablation. Ratios were plotted in GraphPad Prism 9 and compared to normalized contact length before ablation.

#### Correlation analysis between apical cell area and the signal intensity of fluorescent reporters

We measured the mean signal intensity of F-actin (using Utr::GFP), MyoII (using MyoII::mCh), RhoGEF2::GFP, RhoGAP71E::GFP and apical cell area at around 28h APF in time-lapse movies with a time resolution of 5 seconds. Using the Fiji polygon selection tool, we manually traced individual apical perimeters of 2° LCs. ROIs were generated, and minimum, maximum, and mean average signal intensities were obtained for each cell and normalized against the background. Using a custom R script we measured the time-shifted Pearson’s cross-correlations (time windows from ±8 min) between the mean signal intensities of the fluorescent reporter and cell area using 20–30-min-long movies. The data were smoothed using an “R” kernel regression smoother and presented as the average Pearson’s correlation of individual cells and the standard error of the mean. Statistics were performed on the average R-value changes for apical cell area vs. RhoGEF2::GFP, apical cell area vs. RhoGAP71E::GFP, apical cell area vs. Utr::GFP, Utr::GFP and Sqh::mCh, apical cell area vs. MyoII in Act>RhoGAP71E RNAi and RhoGEF2 RNAi expressing cells using the one-sample t-Test with a theoretical mean of 0. Pulse duration was obtained after plotting the apical cell area against time. Distances between the maxima were obtained and averaged. Amplitudes were obtained by calculating the distance between the maxima and minima to obtain an average amplitude or fold-change of signal intensity. For the cell-to-cell comparisons, R-values were generated for the correlation between areas of adjacent 2° LCs and between area of one cell and actin intensity in the neighboring cells. R-values were averaged and a one-sample t-Test was performed against a theoretical mean of 0.

#### Comparing signal intensities between WT and RNAi-expressing cell clones

LCs were selected for analysis if all adjacent cells expressed the RNAi, or, for the control group, if all adjacent cells were WT. For each 2° LCs, circular ROIs were used to measure the signal intensity at the medioapical region from which the background was subtracted to calculate the average signal intensity in RhoGAP71E RNAi or RhoGEF2 RNAi compared with WT LCs. A t-test for normally distributed data and a Mann-Whitney test for non-normally distributed data were used to compare the groups.

#### Data to appear as online supplemental materials

Fig. S1 shows medioapical actomyosin network assembly imaged at a high spatial and temporal resolution that negatively correlates with cell area fluctuations. Fig. S2 shows negatively correlated cytoskeletal and mechanical coupling between anterior and posterior cone cells and 1° cells. Fig. S3 shows that RhoGEF2 overexpression rescues cell shape and rearrangements defects induced by RhoGAP71E overexpression. Movie 1 shows medioapical F-actin network dynamics in WT LCs including ring assembly and node fusion. Movie 2 shows anisometric apical area relaxation of 2° and 3° LCs after laser ablation of their medioapical actomyosin network. Movie 3 shows that Rho1 overexpression accelerates medioapical F-actin dynamics and cell area fluctuations of LCs compared with WT. Movie 4 shows that medioapical actomyosin network assembly and cell area fluctuations are inversely coordinated between neighboring 2° LCs. Movie 5 shows that constitutive F-actin and MyoII assembly disrupt medioapical actomyosin organization and dynamics. Movie 6 shows that RhoGEF2 promotes medioapical F-actin and MyoII dynamics in LCs. Movie 7 shows that RhoGAP71E inhibits medioapical F-actin and MyoII dynamics in LCs.

## SUPPLEMENTAL FIGURE LEGENDS

**Figure S1:**
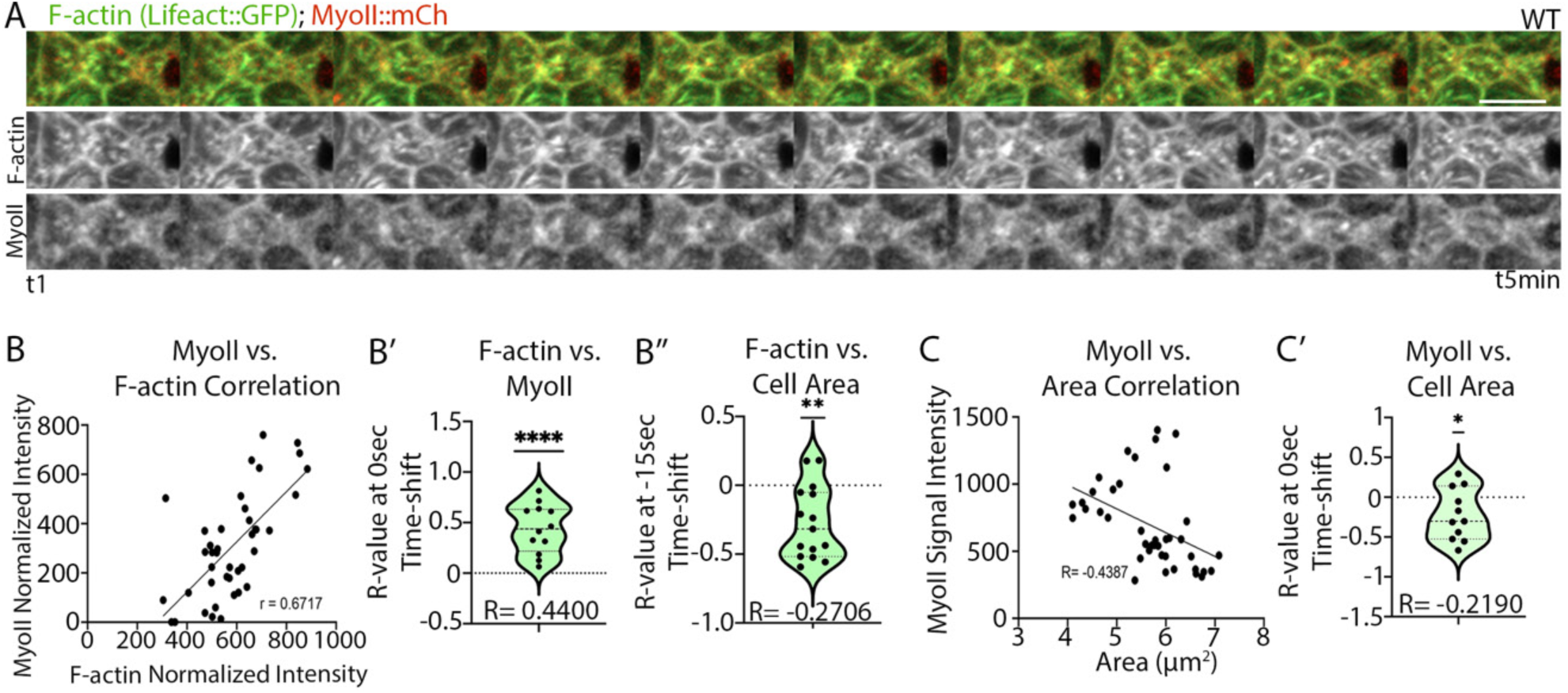
Pulsatile medioapical actomyosin network assembly correlates with apical cell area contractions. (A) High-resolution Airyscan time-lapse movie of F-actin (Lifeact::GFP) and MyoII (MyoII::mCh) taken every 30 sec. Note F-actin and MyoII accumulation in nodes and in filamentous structures. (B) Positive Pearson’s correlation between F-actin and MyoII in a single LC. (B’) Time-shifted Pearson’s correlation between medioapical F-actin and MyoII shows a strong positive correlation at 0 sec time-shift (mean R= 0.4400, N=15 cells, one-sample t-Test: p<0.0001). (B’’) Time-shifted Pearson’s correlation between medioapical F-actin and apical cell area shows a negative correlation at a −15 sec time-shift (mean R= −0.2706, N=15 cells, one-sample t-Test: p=0.0014). (C) Negative Pearson’s correlation between MyoII intensity and apical cell area in a single LC. (C’) One sample t-Test shows a significant negative correlation between medioapical MyoII levels and cell area at a 0 sec time-shift (mean R= −0.2190, N= 11, p=0.0482). Scale bar in this and subsequent figures is 5 µm.

**Figure S2:**
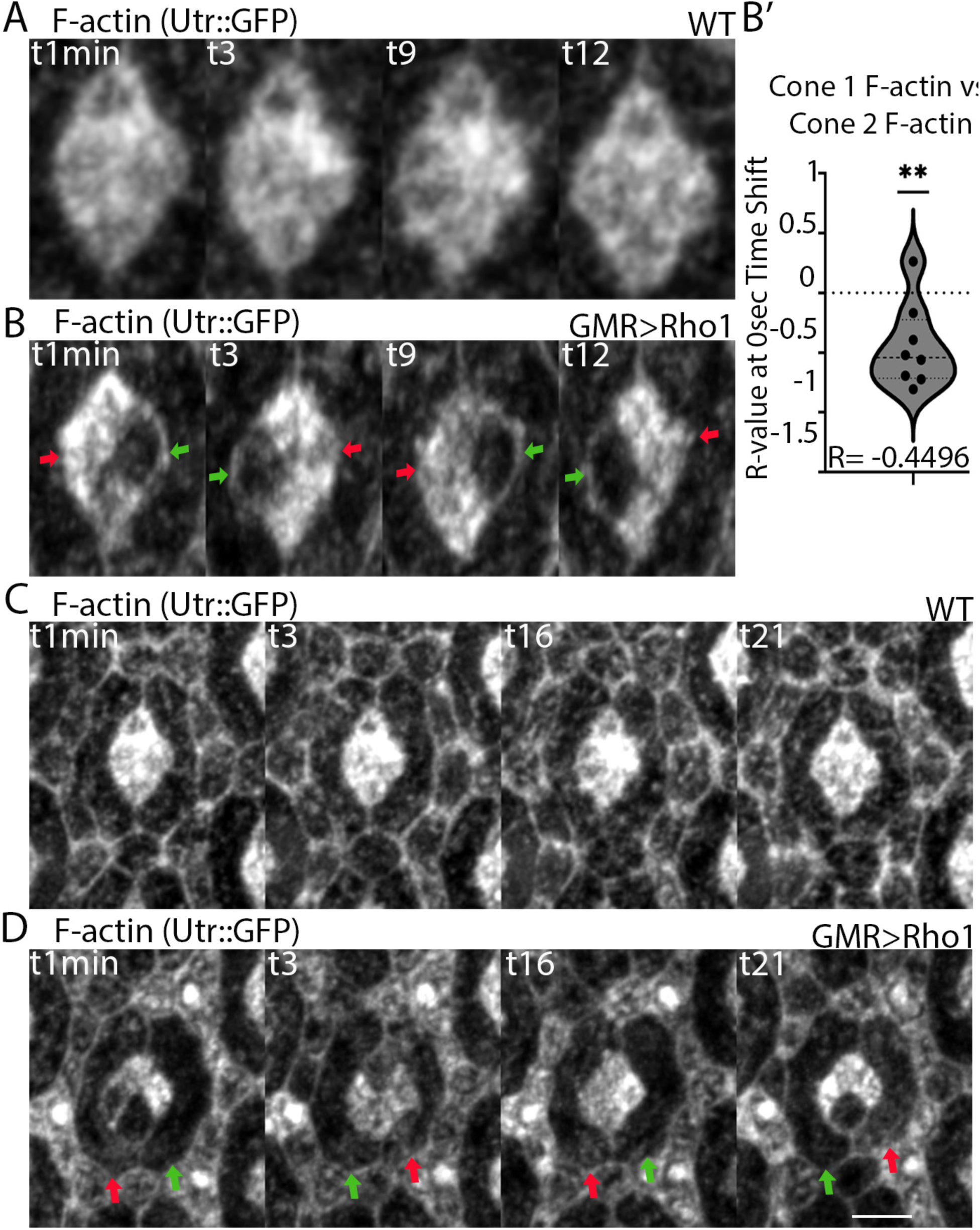
Anterior and posterior cone cells and 1° cells are mechanically coupled. (A-D) Rho1 overexpression promotes strong fluctuations in F-actin levels and apical cell area fluctuations that are inversely coordinated between neighboring cells. (A-B) Kymograph showing typical fluctuations in medioapical F-actin and cell area in (A) WT cone cells and (B) cone cells that overexpress Rho1. Green arrows, low F-actin intensity and apical cell area relaxation. Red arrows, high F-actin intensity and apical cell area contraction. (B’) Fluctuations in F-actin levels are negatively correlated between anterior and posterior cone cells that overexpress Rho1 (R=−0.4496, N=8 pairs, 16 cells, one-sample t-Test: p=0.0086). (C-D) Kymographs of medioapical F-actin dynamics and apical cell area fluctuations in (C) WT and (D) Rho1-overexpressing 1° LCs. (D) Fluctuations in F-actin and apical cell area also appear negatively correlated between adjacent 1° LCs expressing Rho1.

**Figure S3:**
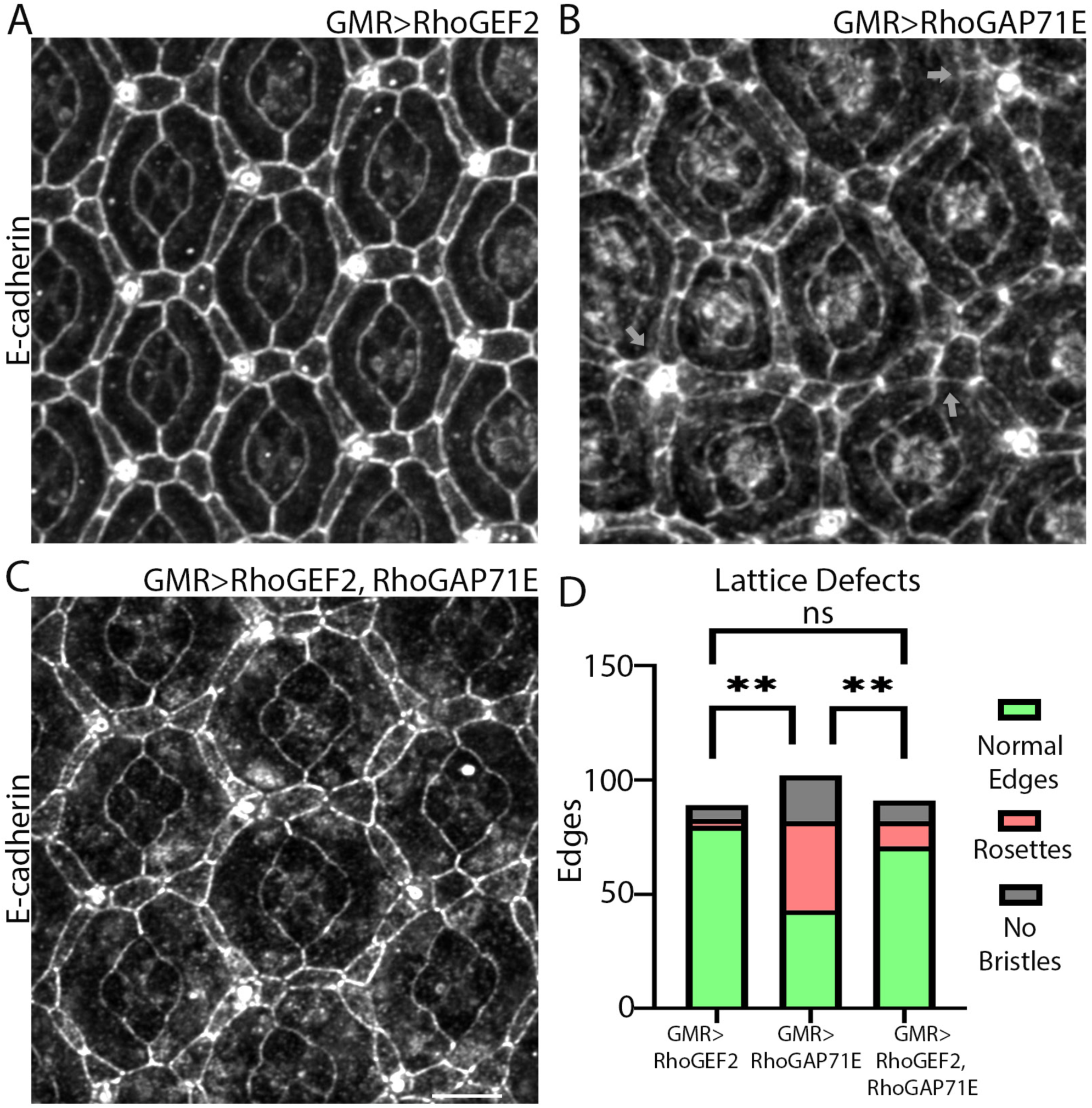
RhoGEF2 rescues cellular defect caused by RhoGAP71E expression. (A) Epithelial organization in GMR>RhoGEF2-expressing eyes where epithelial cells maintain normal shape and rearrangements. (B) Expressing GMR>RhoGAP71E induces intercalation defects and expansion of apical cell area. (C) Expressing RhoGEF2 in RhoGAP71E-expressing eyes partially rescued the GMR>RhoGAP71E phenotypes. (D) Quantification of lattice defects. One-way ANOVA with Tukey’s multiple comparisons comparing the mean of changes in formation of rosettes. RhoGEF2 vs. RhoGAP71E, p=0.0015. RhoGAP71E vs. RhoGEF2/RhoGAP71E, p=0.0056. RhoGEF2 vs. RhoGEF2/RhoGAP71E, p=0.3789, N= 80-100 edges.

## SUPPLEMENTAL MOVIES

**Movie 1: Medioapical F-actin network dynamics in WT LCs.** (Corresponds to Fig. 1). Time-lapse videos of F-actin (Utr::GFP) in WT that highlight medioapical actomyosin dynamics. Left panel: Yellow arrowheads show the fusion of two F-actin nodes in the 2° LC demarcated by the white box. Right panel: Red arrow shows the formation of a medioapical ring-like structure, the subsequent disassembly of the ring and actin flow to the LC-LC contact (note the green arrow movement). Scale bar in this and subsequent movies is 5 µm.

**Movie 2: Medioapical actomyosin network ablation induces asymmetric cell expansion of the 2° and 3° LCs.** (Correspond to Fig. 2). Time-lapse videos of a 2° (left panel) and a 3° (right panel) LCs with tagged F-actin (Utr::GFP) and MyoII (MyoII::mCh). Targeted ablations of the medioapical actomyosin network in the LCs induced a rapid anisometric apical cell area relaxation (note the area of the 2° and 3° LCs highlighted by the white box before and after ablation).

**Movie 3: Rho1 expression accelerates medioapical F-actin dynamics and cell area fluctuations in LCs.** (Corresponds to Fig. 3). Time-lapse videos of F-actin (Utr::GFP) dynamics in WT retina (left panel) and in a retina that overexpresses Rho1 (right panel). Note the robust and accelerated assembly and disassembly of medioapical F-actin in the LCs. This faster turnover of the medioapical actomyosin network accompanied a rapid decrease and increase in the apical cell area. Red and green arrowheads highlight the assembly and disassembly of the medioapical actomyosin network, respectively.

**Movie 4: Medioapical actomyosin network assembly and cell area fluctuations are inversely coordinated between adjacent 2° LCs.** (Corresponds to Fig. 3). Time-lapse video of a fly retina that overexpresses Rho1 compared with WT. Rho1 overexpression accelerated the assembly and disassembly of the medioapical F-actin networks and cell area contraction and expansion. Note that this behavior is inversely coordinated between the two neighboring 2° LCs demarcated by the white box. The green arrowhead marks medioapical F-actin assembly; red arrowhead marks disassembly.

**Movie 5: Constitutive F-actin and MyoII assembly disrupts medioapical actomyosin organization and dynamics.** (Corresponds to Fig. 4). (Left) Time-lapse video of MyoII (MyoII::mCh) in a MLCK^CA^-expressing eye. Note the rapid assembly and disassembly of a medioapical ring marked with yellow arrowheads. (Right) A time-lapse video of F-actin (Utr::GFP) in a Dia^CA^-expressing eye. Note the increased F-actin levels around the LCs’ borders and the low levels and dynamics of medioapical F-actin.

**Movie 6: RhoGEF2 accelerates medioapical F-actin and MyoII dynamics in LCs.** (Corresponds to Fig. 5) Left: A time-lapse video of F-actin (Utr::GFP) in an eye overexpressing RhoGEF2. Note the rapid assembly and disassembly of the medioapical F-actin network marked by the red and yellow arrowheads, respectively. Middle and right: A time-lapse video of MyoII (MyoII::mCh) in clone cells that express a RhoGEF2 RNAi (GFP^+^, white arrowhead) compared with WT counterparts (yellow arrowhead). Right: Note loss of MyoII intensity in the RhoGEF2 RNAi-expressing 2° LC compared with the WT.

**Movie 7: RhoGAP71E inhibits medioapical F-actin and MyoII dynamics in LCs.** (Correspond to Fig. 6) Left: Time-lapse video of F-actin (Utr::GFP) in eyes that overexpress RhoGAP71E. Note the increased apical cell area of the LCs and reduced medioapical F-actin levels. Middle: Time-lapse video of MyoII (MyoII::mCh) in clones expressing RhoGAP71E RNAi, positively marked with RFP. Right: Note the reduced apical cell area of the RhoGAP71E RNAi expressing cell (white arrowhead) compared with WT (yellow arrowhead).

